# Reaction-Diffusion Mechanisms Underlying Hirschprung’s Disease and their Practical Impications

**DOI:** 10.1101/492884

**Authors:** Arturo Tozzi

## Abstract

The erratic extent of aganglionic and hypoganglionic segments in Hirschsprung’s disease (HD) makes it difficult to predict the amount of the intestine to remove in order to restore the proper functional motility. Our aim was to assess whether the embryonic rostro-caudal intestinal colonization by neuroblasts from the neural crest follows a predictable pattern in HD. In touch with Turing’s reaction diffusion model (RD), which describes biological patterns (such as leopard spots and lung branching morphogenesis) in terms of interactions/competitions between activating and inhibiting factors, we hypothesized that intestinal neural density could be triggered by local gut factors that counteract the proximal-distal embryonic progression of neural progenitors. While the neuronal number is approximately the same throughout the whole intestine in healthy subjects, in HD neural density decreases rostro-caudally towards the rectal region, due to an augmented activity and concentration of distal local inhibitors. In order to prove our hypothesis of HD’s nervous rostro-caudal adjustments driven by Turing-like processes, we compared the neuronal density patterns achieved through RD models’ simulations with the neuronal numbers detected in different colonic regions from affected children. We showed that the virtual and the real plots display fully overlapping and matching features. The fact that neuronal decreases in impaired colons match Turing equations’s previsions points towards the human intestine (both healthy and sick) as colonized through a diffusive proximal-distal neural pattern that is predictable, allowing us to straightforwardly calculate the length of the gut to resect during surgical procedures for HD.

---

During prenatal embryonic development, cells from the neural crest migrate rostro-caudally into intestine to form the myenteric and submucosal plexuses. In Hirschsprung’s disease (HD) (Hirschsprung 1888) neurons are missing from the distal parts of the intestine, giving rise to aganglionic and hypoganglionic transitional zones of variable extent (Han et al., 2018). The extremely variable length of the impaired intestine is a painstaking problem that occasionally prevents surgeons to fully remove the pathological gut segment and restore the proper intestinal motility (Miele et al., 2000; Genser et al., 2013). Here we ask: does the pathological intestinal colonization in HD follow a predictable pattern? Is it feasible to calculate the length of the gut that must be surgically removed, in order to avoid to leave inside the abdomen hypoganglionic segments, which prevent functional recovery of gut motility?

Here Turing’s reaction-diffusion model (RD) comes into play (Turing 1952). In its early formulation, RD describes a system of chemical substances, where random disturbances occur due to the competition of two active components, termed activators and inhibitors (Deca, 2017). The balance between excitatory and inhibitory inputs with different diffusion coefficients gives rise to peculiar diffusion patterns of the chemical substances into the system’s lattice (Kondo and Miura 2010). In the last sixty years, RD has been proven able to describe different biological dynamics and features, such as, for example, zebrafish markings, shark denticles, avian feathers, leopard spots and grid cell’s firing patterns in the brain (Kondo et al., 2009; Cooper et al., 2018).

Here we hypothesize that an RD mechanism might explain the lack of neuronal colonization of the proximal gut in case of HD, giving rise to aganglionic and hypoganglionic areas where the physiological neural density is impaired. Our aim is to introduce a model of nervous colonization of the gut where reagents (standing for neuroblasts from the neural crest) enter a cylinder (i.e., the enteric wall) from an extremity (i.e., the proximal intestine where the neural progenitors enter the gut during embryonic development) and homogeneously and ergodically diffuse towards the other extremity (i.e., the distal gut towards the more caudal rectal segment). While the neural density is preserved throughout the whole normal gut, the neuronal number in HD, higher in the proximal intestine, progressively decreases towards the distal areas, due to local inhibition factors which counteract the proper rostro-caudal colonization from neural progenitors. Here we provide the proper Turing equations and compare the theoretical results from RD simulations with the available experimental data from real neural concentration in the impaired colon of children affected by HD. The matching of our theoretical results and real data would be able to provide surgeons with an central information: the exact proximal point where to cut the gut in order to fully remove the affected intestine. In sum, starting from the neuronal density in different colonic areas of HD patients, we aim to prove that embryonic neuronal intestinal migration (in both healthy and sick individuals) follows Turing’s RD equations in a way that is predictable. Therefore, if we are able to correlate theoretical RD models with the real patterns of neural colonization, we achieve two goals: we demonstrate that the gut is colonized through a diffusion pattern of nervous structures, counteracted by local factors endowed in the distal gut; further, we accurately estimate the length of the pathological intestine to remove during surgical procedures for HD.

## MATERIALS AND METHODS

### Theoretical framework

Following Turing (1952), we consider partial differential equations representing the number of perturbations (spatial variations) occurring in the system under evaluation, i.e., the embryonal human intestine colonized by neural precursors originating from the neural crest. RD modeling of the proximal-distal neuronal colonization of the gut is undertaken by using a modification of the Turing’s standard activator-inhibitor model (Kondo and Miura, 2009). The required master equations are (Cooper et al., 2018):

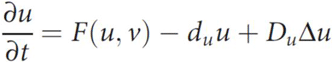

And:

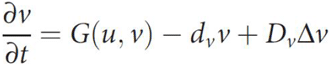

Where u stands for the concentration of the neuronal progenitors diffusing from the proximal towards the distal intestine (activator) and v for the concentration of the counteracting force inside the gut (an inhibitory force, that is stronger in the distal than in the proximal intestine), which prevents the proper colonization achieved by u. In sum, the above-mentioned equations describe the rate change of the neuronal number (concentrations) in time and space, due to the diffusion of u partially contrasted by v.

The term F(u,v) stands for a nonlinear function defined by (Cooper et al., 2018):

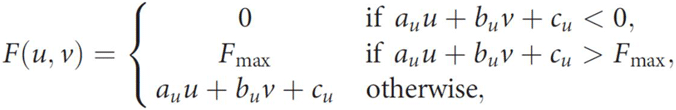

While the term G(u,v) stands for another nonlinear function, defined by:

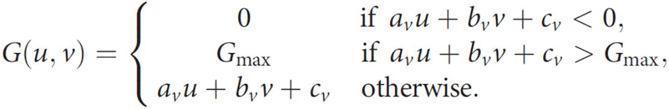

A simulation may be generated by Phyton code, taking into account that the prescribed initial conditions and starting parameters may vary across different simulations (**Figure 1A**). Indeed, a fine-tuning of several parameters may be accurately performed: for example, by exactly defining the center of a spot in an initiator row of a given number and radius; by choosing a spatial discretization and a sufficient small-time step to ensure numerical stability; and by changing the concentrations of the two active reagents u and v (the activator and the inhibitor, respectively). **Figure 1A** illustrates the general sketch and a feasible simulation. For technical readers, the required equations, parameters and mathematical steps are described in: Cooper et al. (2018); Xu et al. (2017).

In terms of the physiological intestinal rostro-caudal colonization by neural crest’s neural precursors (**Figure 1B**, **left side**), this means that unchanged concentrations of activator lead to a constant number of neurons throughout the whole gut. In turn, in HD, the decrease of the activator and/or the increase of the inhibitor lead to an anomalous neuronal diffusion, with a progressive proximo-distal decrease in nervous density (**Figure 1B**, **right side**).

### Comparison with human data

The above-mentioned changes in neuronal concentration occurring in the dynamical process of rostro-caudal diffusion of neuroblast are predictable: in the Results, we will provide the curves and paths described by the RD model, comparing them with real human data. Concerning the latter, we retrospectively evaluated the results from one of the best available published paper, i.e., Osterheld et al. (2009). These Authors used standardized methodology for the examination of resected bowel after Duhamel pull-through procedure for HD. Following the histological standardized technique described by Meier-Ruge in children (Meier-Ruge et al., 1999), Osterheld et al. (2009) used HE and Hematoxylin-Phloxin-Safran stainings and acetylcholinesterase histochemistry on “swiss roll” samples. They analyzed the whole length of resected colons from six patients with HSCR with age at surgery between 1,5 months and 6 years. They looked for a wide range of quantifiable parameters: length of the resected colon, proximal/distal diameters of the removed specimen, number of ganglia/2cm in submucosal and myenteric plexuses, number of cells/2cm in submucosal and myenteric plexuses, nerve hypertrophy in submucosal and myenteric plexuses, AchE activity in submucosal plexus.

**Figure 1A:**
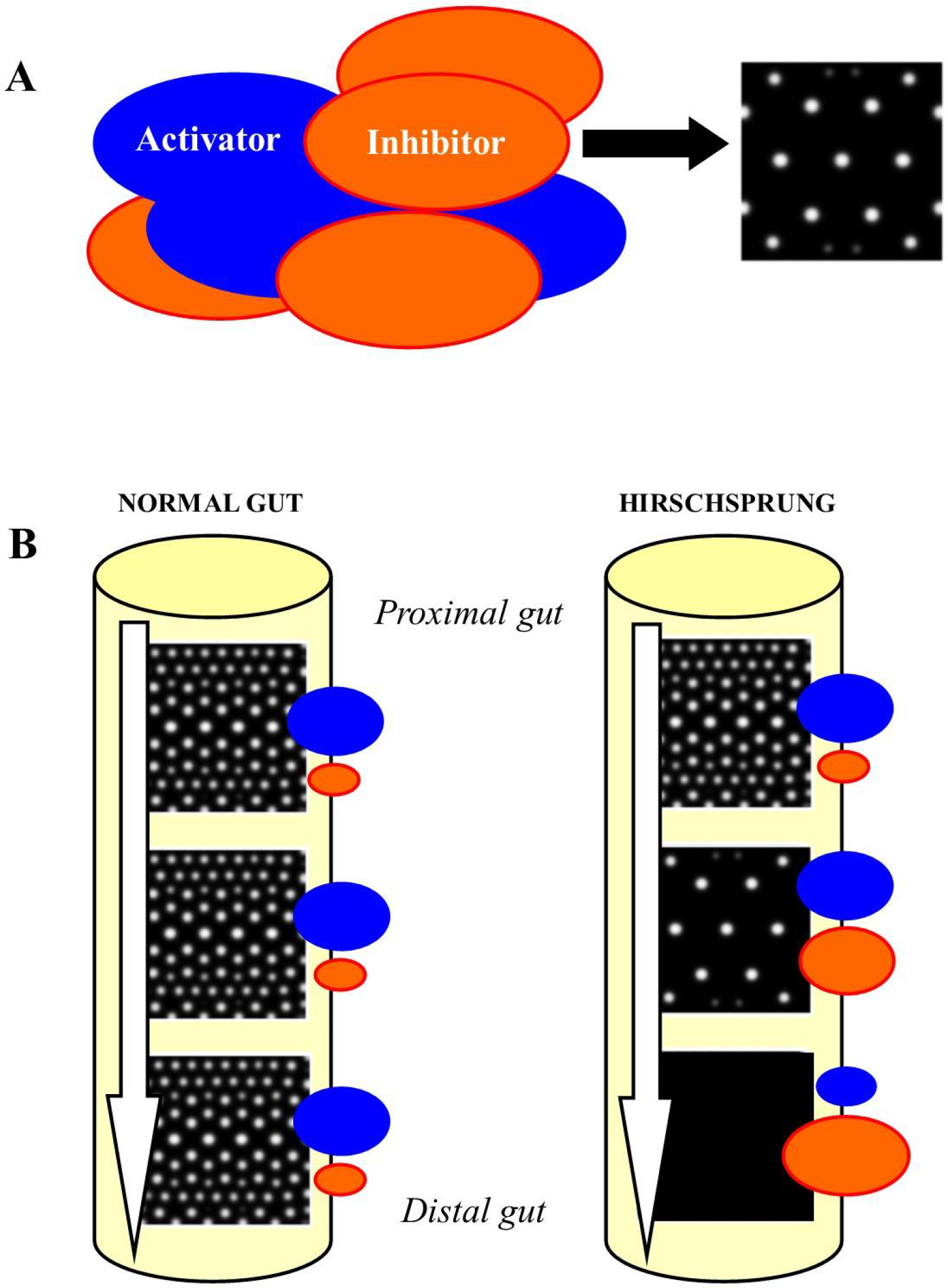
Simulation of Turing’s spots production, starting from the proper initial parameters. The blue circles stand for the activators’ concentration, the red ones for the inhibitor’s concentration. The interaction between the two biological factors gives rise to a spotted pattern on a two-dimensional lattice. In neural terms, the white spots in the right picture stand for the neural density which can be detected in a full-thickness intestinal layer stained with H&E. **Figure 1B**: In the healthy gut, embryonal progressive colonization by neuronal progenitors from neural crest takes place (**Figure 1B**, **left side**). This path follows a proximal-distal (rostro-caudal) trajectory, so that the final number of neurons will be approximately the same in all the segments of the adult gut. In case of HD (**Figure 1B**, **right side**), the number of neurons decreases during the proximo-distal colonization. In Turing models’ terms, the colonization path (white arrow) is counteracted and locally blocked by inhibitory factors displaying higher concentrations in the distal gut. In HD, due to slight changes in the initial conditions of different zones, the spots are allowed to decrease and then to disappear: this leads to hypo-/aganglionic pathological segments.

## RESULTS

The diffusive paths and changes in neuronal concentration described by RD models and their master equations are predictable. Simulations through Pyton give rise to the following results: in the healthy intestine, the nervous density pattern on a plot can be described by a straight line standing for a preserved, constant number of neurons throughout the whole gut (**Figure 2A**). In turn, in HD, RD equations produce a curved line that follows a logistic curve (**Figure 2A**). When comparing the above-mentioned theoretical curves from RD simulations with the ones extracted from real data, we notice a remarkable superimposition between the two paths. Indeed, the submucosal neuronal density in patients affected by HD which underwent pull-through surgical procedure for removal of aganglionic/hypoganglionic segments (**Figures 2A and 2B**), follows the same logistic curve produced by the above-mentioned simulations (**Figure 2B**). Note that we choose, in touch with Osterheld et al. (2009), to mainly focus on the number of submucosal neurons/2cm of colon. Nevertheless, the plots extracted by other data (such as the number of myenteric neurons/2cm of colon, the number of ganglia, and so on) fully overlap the curve of the submucosal plot described in **Figure 2B**.

It is noteworthy that, at a given cutoff value of submucosal neuronal density, the slope takes a steep ascend. This means that the maximum density value, corresponding to the normoganglionic intestine, can be found slightly above the cutoff value (**Figure 2A**). If we consider **Figure 2B**, we notice that, at the cutoff value of about 50-60 submucosal neurons, we are almost certain to find the normoganglionic gut located 4-5 centimeters proximally. This means that, during a surgical procedure for HD, when the surgeon detects about 55-60 submucosal neurons in intraoperative samples stained with H&E, he is allowed to safely cut five centimeters above: he will be sure to avoid the risk to leave hypoganglionic segments inside the abdomen.

**Figure 2A:**
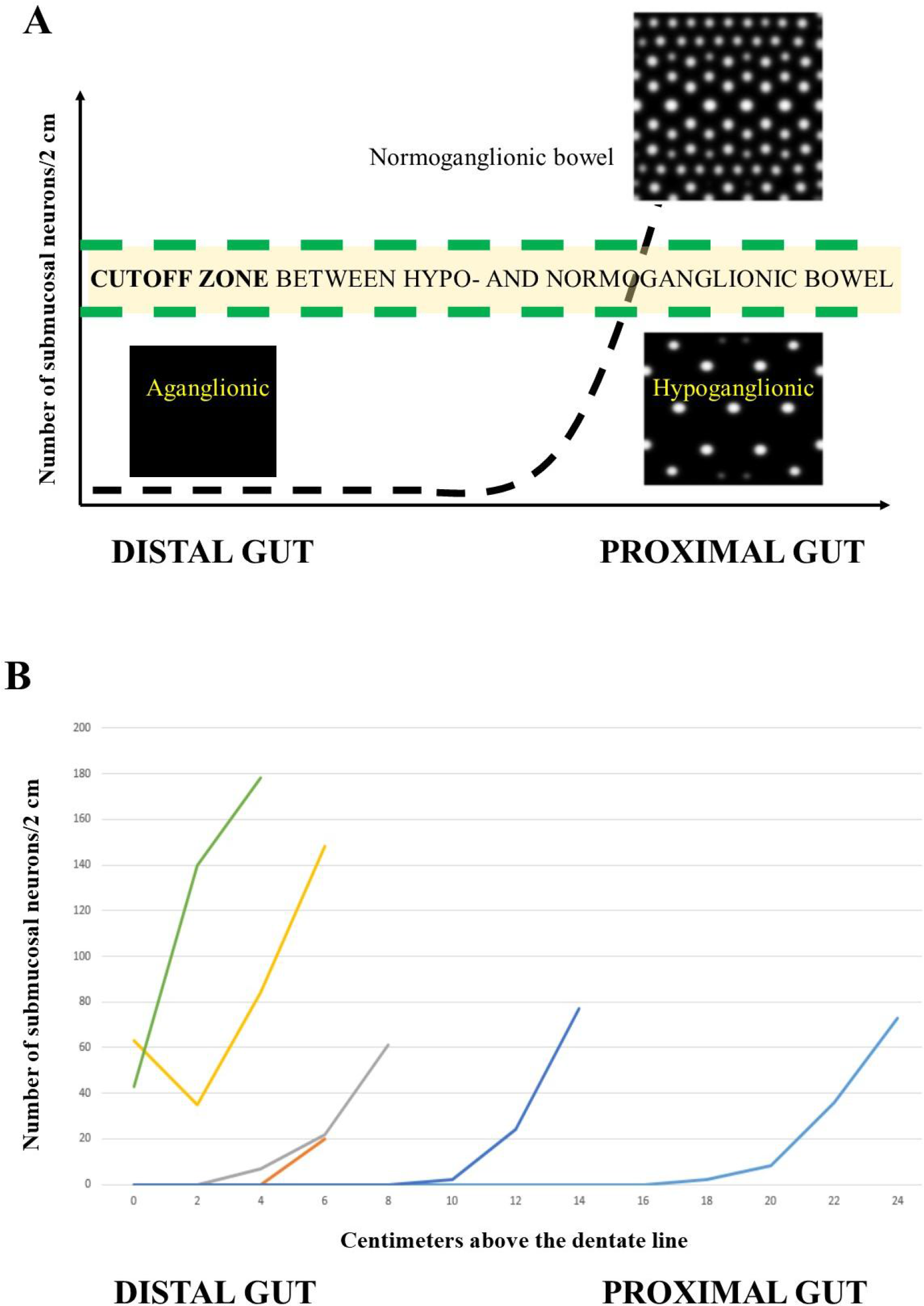
Interactions between activation and inhibition are quantifiable in terms of Turing equations. In a normal intestine, the number of neurons colonizing the gut will remain the same in all the zones (green upper dotted straight line). In turn, the number of neurons will be reduced in HD, following the path of a logistic curve (black dotted curved line). The Figure also illustrates a zone of cutoff values between the healthy and the diseased colon: for this crucial issue, see the main text. **Figure 2B**: submucosal neuronal density from colons of six patients surgically treated for HD. The plot summarizes the number of neurons found in the six resected segments at different lengths from the dentate line (the zero value on the x axis stands for the dentate line, close to the anus). It is easy to notice that a decrease occurs in the number of submucosal ganglion cells, from the proximal to the distal parts of the resected colons. Data extrapolated from Osterheld et al., 2009.

## DISCUSSION

We found that the density pattern of neurons predicted by RD models’ simulations match the experimental findings detected in the intestine of patients affected by HD. Here we wanted to test the hypothesis that craniocaudal migration of neuroblasts can be described in terms of Reaction-Diffusion systems. The solutions of RD equations describe a wide range of behaviors, including the formation of travelling waves and wave-like phenomena, as well as other self-organized patterns like stripes, hexagons or the more intricate structures of dissipative solitons. Such patterns have been dubbed “Turing patterns” and are able to describe, for example, not just the spots of leopards or the stripes of zebras, but also more complex phenomena such as lung branching morphogenesis (Xu et al., 2017). The most accepted theory states that the HD’s embryonic disorder in caused by a lack of craniocaudal migration, differentiation and maturation of neuroblasts from the neural crests, taking place by the fifth to the twelfth week of human gestation: the earlier the migration ceases, the longer the aganglionic segment will be, leading to more or less severe constipation. Defects in the differentiation of neuroblasts into ganglion cells and accelerated ganglion cell destruction within the intestine may also contribute to the disorder. For further details, see: Genetics Home Reference, https://ghr.nlm.nih.gov/condition/hirschsprung-disease. The rationale of using an RD model to describe HD pathogenetic mechanisms lies in experimental data that seem to integrate the standard, canonical account. Indeed, other causes might theoretically contribute to the incomplete migration of neural progenitors in HD. Landman et al. (2003) suggested that, during embryonic development, cell migrations takes place on an underlying tissue domain which is itself growing over the time scale of days in which this phenomenon occurs. Once established that cell migration/colonization is strongly affected by the tissue domain growth (as it occurs in the embryonic gut), the Authors provided mathematical approximation to the full system equations, by using a simplified chemotactic migration model which closely resembles RD models: in particular, they considered either linear, exponential and, above all, the logistic uniform domain growths which are in touch with our account.

HD can result from mutations in one of several genes, including, among others, the RET, EDNRB, EDN3 and semaphorins genes (Auricchio et al., 1999; Tang 2012; Jiang et al., 2015). However, the genetics of this condition appear complex and are not completely understood. While a mutation in a single gene sometimes causes the condition, mutations in multiple genes may be required in some cases (Alves et al., 2013). Indeed, the genetic cause of HD is unknown in half of affected individuals. Further, many of the above-mentioned involved genes, apart from playing an important and well-established role in the formation of normal enteric nerves, are also linked with mesenchymal activities which could be expressed in the distal colon. In touch with this observation, an RD model of HD points towards the possibility of local factors located in the distal gut’s environment able to inhibit the canonical craniocaudal progression of neuroblasts. Here we provide a few examples which demonstrate that, in touch with our account of a strict interaction between RD’s activating and inhibitory active factors in the distal gut, the cross-relationships among different genes products may give rise to disturbances in the balance of rostro-caudal neuroblastic colonization of the intestine. Bondurand et al. (2000) showed that PAX3 and SOX10 interact directly by binding to a proximal region of the MITF promoter, leading. in the dominant megacolon mouse, to impaired MITF expression as well as melanocytic development and survival. Furthermore, Lee et al. (2000) demonstrated that wildtype SOX10 directly binds and activates transcription of the MITF promoter, whereas a mutant form of the SOX10 protein associated with Waardenburg-Shah syndrome acts as a dominant-negative repressor of MITF expression and reduces endogenous MITF protein levels. Bondurand et al. (2001) showed that SOX10, in synergy with EGR2, strongly activates Connexin-32 expression in vitro by directly binding to its promoter. In agreement with these findings, SOX 10 and EGR2 mutants in patients with peripheral myelin defects failed to transactivate the CX32 promoter. In addition, some CMTX1 patients harbored a T-to-G transversion at position −528 of the CX32 promoter. The authors demonstrated that this mutation eliminates binding and activation by SOX10. By assessing expression of Sox9 or Sox10 in early Xenopus embryos, Taylor and LaBonne (2005) demonstrated that each one of these factors could direct the formation of neural crest precursors and the development of a range of neural crest derivatives. Also, they identified Sumo1 and Ubc9 as Sox-interacting proteins that play a role in regulating the function of Sox9 and Sox10 during neural crest and inner ear development.

An RD model for intestinal migration of neuroblasts predicts that the subtle balance between activators’ and inhibitors’ concentration gives rise to aganglionic, hypoganglionic or normoganglionic gut segments. However, a further, sometimes unnoticed option must be taken into account: an hyperganglionic intestine, morphologically characterized by hyperplasia of submucosal and myenteric plexuses (Tozzi et al., 1999). In RD terms, hyperganglionosis might stand for the absence of the slight amount of the distal inhibitor (**see Figure 1B, left side**) required for the normal development of healthy gut. HD may be associated with other conditions, such as Waardenburg syndrome, which display pigmentation changes resembling a Turing-like mechanism. In touch with this latter claim, an animal counterpart of human aganglionic megacolon comes into play: the lethal white foal syndrome, which is a congenital abnormality of overo spotted horses. These foals display, apart an intestinal obstruction that proves fatal within the first few days of life, a nearly all-white coat similar to Turing spots.

RD models, apart from providing novel insights in HD pathogenetical mechanisms, also suggest real-world consequences concerning the proper technical approaches to the required surgical procedures. Treatment of HD is generally by surgery to remove the affected section of bowel, including the (more distal) aganglionic and the (more proximal) hypoganglionic segments. After removal of the impaired segment, a pull-through is performed in order to link the normally ganglionic bowel with the anus. However, the length of the affected bowel is extremely variable, ranging from few colonic centimeters above the dentate line to several meters, in the forms involving a variable amount of the short bowel too. Therefore, it is from time to time very difficult to establish how much intestine needs to be removed, because both the aganglionic and hypoganglionic areas display high interindividual variability. In order to tackle this practical issue, we showed that RD methodology is able to accurately predict, by using an unpretentious H&E staining in intraoperative biopsies, how many centimeters of colon need to be removed to prevent uncomplete surgical bowel resection.

## REFERENCES

1) Alves MM, Sribudiani Y, Brouwer RW, Amiel J, Antiñolo G, et al. 2013. Contribution of rare and common variants determine complex diseases-Hirschsprung disease as a model. Dev Biol. 2013 Oct 1;382(1):320–9. doi: 10.1016/j.ydbio.2013.05.019. Epub 2013 May 23. Review.

2) Auricchio A, Griseri P, Carpentieri ML, Betsos N, Staiano A, Tozzi A, Priolo M, Thompson H, Bocciardi R, Romeo G, Ballabio A, Ceccherini I. 1999. Double Heterozygosity for a RET Substitution Interfering with Splicing and an EDNRB Missense Mutation in Hirschsprung Disease. The American Journal of Human Genetics 05/1999; 64(4):1216–21., DOI:10.1086/302329.

3) Bondurand N, Pingault V, Goerich DE, Lemort N, Sock E, et al. 2000. Interaction among SOX10, PAX3 and MITF, three genes altered in Waardenburg syndrome. Hum Mol Genet. 2000 Aug 12;9(13):1907–17.

4) Bondurand N, Girard M, Pingault V, Lemort N, Dubourg O, Goossens M. 2001. Human Connexin 32, a gap junction protein altered in the X-linked form of Charcot-Marie-Tooth disease, is directly regulated by the transcription factor SOX10. Hum Mol Genet. 2001 Nov 15;10(24):2783–95.

5) Cooper RL, Thiery AP, Fletcher AG, Delbarre DJ, Rasch LJ, Fraser GJ. 2018. An ancient Turing-like patterning mechanism regulates skin denticle development in sharks. Sci Adv. 2018 Nov 7;4(11):eaau5484. doi: 10.1126/sciadv.aau5484. eCollection 2018 Nov.

6) Deca D. 2017. Grid Cells-From Data Acquisition to Hardware Implementation: A Model for Connectome-Oriented Neuroscience. In: The Physics of the Mind and Brain Disorders: Integrated Neural Circuits Supporting the Emergence of Mind, edited by Opris J and Casanova MF. New York, Springer; Series in Cognitive and Neural Systems. Pages 493–511. ISBN: 978-3-319-29674-6. DOI10.1007/978-3-319-29674-6_9.

7) Genser L, Manceau G, Karoui M, Breton S, Brevart C, et al. 2013. Postoperative and long-term outcomes after redo surgery for failed colorectal or coloanal anastomosis: retrospective analysis of 50 patients and review of the literature. Dis Colon Rectum. 2013 Jun;56(6):747–55. doi: 10.1097/DCR.0b013e3182853c44.

8) Han JW, Youn JK, Oh C, Kim HY, Jung SE, Park KW. 2018. Why Do the Patients with Hirschsprung Disease Get Redo Pull-Through Operation? Eur J Pediatr Surg. 2018 Aug 1. doi: 10.1055/s-0038-1667038.

9) Hirschsprung H. 1888. Stuhlträgheit Neugeborener in Folge von Dilatation und Hypertrophie des Colons. Jahrbuch für Kinderheilkunde und physische Erziehung. Berlin. 27: 1–7.

10) Jiang Q, Arnold S, Heanue T, Kilambi KP, Doan B, et al. 2015. Functional loss of semaphorin 3C and/or semaphorin 3D and their epistatic interaction with ret are critical to Hirschsprung disease liability. Am J Hum Genet. 2015 Apr 2;96(4):581–96. doi: 10.1016/j.ajhg.2015.02.014.

11) Kondo S, Iwashita M, Yamaguchi M. 2009. How animals get their skin patterns: fish pigment pattern as a live Turing wave. Int J Dev Biol. 2009;53(5-6):851–6. doi: 10.1387/ijdb.072502sk.

12) Kondo S, Miura T. 2010. Reaction-diffusion model as a framework for understanding biological pattern formation. Science. 2010 Sep 24;329(5999):1616–20. doi: 10.1126/science.1179047.

13) Landman KA, Pettet GJ, Newgreen DF. 2003. Mathematical models of cell colonization of uniformly growing domains. Bull Math Biol. 2003 Mar;65(2):235–62.

14) Lee M, Goodall J, Verastegui C, Ballotti R, Goding CR. 2000. Direct regulation of the Microphthalmia promoter by Sox10 links Waardenburg-Shah syndrome (WS4)-associated hypopigmentation and deafness to WS2. J Biol Chem. 2000 Dec 1;275(48):37978–83.

15) McCabe L, Griffin LD, Kinzer A, Chandler M, Beckwith JB, McCabe ER. 1990. Overo lethal white foal syndrome: equine model of aganglionic megacolon (Hirschsprung disease). Am J Med Genet. 1990 Jul;36(3):336–40.

16) Meier-Ruge WA, Brunner LA, Engert J, Heminghaus M, Holschneider AM,et al. 1999. A correlative morphometric and clinical investigation of hypoganglionosis of the colon in children. Eur J Pediatr Surg. 1999 Apr;9(2):67–74.

17) Miele E, Tozzi A, Staiano A, Toraldo C, Esposito C, Clouse RE. 2000. Persistence of abnormal gastrointestinal motility after operation for Hirschsprung’s disease. The American Journal of Gastroenterology 06/2000; 95(5):1226–30., DOI: 10.1111/j.1572-0241.2000.02014.x.

18) Xu H Sun M, Zhao X. 2017. Turing mechanism underlying a branching model for lung morphogenesis. PLoS One. 2017 Apr 4;12(4):e0174946. doi: 10.1371/journal.pone.0174946. eCollection 2017.

19) Osterheld MC, Meagher-Villemure K, Ciola AM, Martin P, Vilas D, Meyrat BJ. 2009. Hirschsprung’s disease: the “Swiss roll” technique revisited. Pediatr Surg Int. 2009 Jul;25(7):573–8. doi: 10.1007/s00383-009-2395-x. Epub 2009 Jun 12.

20) Tang C. 2012. Genome-wide copy number analysis uncovers a new HSCR gene: NRG3. PLoS Genet. 8 (5): e1002687. doi:10.1371/journal.pgen.1002687.

21) Taylor KM, Labonne C. 2005. SoxE factors function equivalently during neural crest and inner ear development and their activity is regulated by SUMOylation. Dev Cell. 2005 Nov;9(5):593–603.

22) Tozzi A, Staiano A, Tramontano A, Miele E, Toraldo C. 1999. Hyperganglionosis and Hirschsprung’s disease. Journal of Pediatric Gastroenterology and Nutrition 05/1999; 28(5)., DOI: 10.1097/00005176-19990500000132.

23) Turing AM. 1952. The chemical basis of morphogenesis. Philos Trans R Soc B; Biol Sci 237(641): 37–73.

